# Stereocilium height changes can account for the calcium dependence of the outer-hair-cell bundle’s resting state

**DOI:** 10.1101/2024.11.18.624097

**Authors:** Rayan Chatterjee, Dáibhid Ó Maoiléidigh

## Abstract

Outer-hair-cell bundles are sensory organelles required for normal hearing in mammals. These bundles convert sound-induced forces into receptor currents. This conversion depends on the resting receptor current of each bundle, which increases when extracellular calcium is decreased to the physiological level. How extracellular calcium regulates the bundle’s resting state is not well understood. We propose a mechanism explaining how extracellular calcium can regulate the outer-hair-cell bundle’s resting state. Each bundle comprises filamentous stereocilia linked by gating springs that are attached to ion channels. Sound-induced forces deflect stereocilia, increasing and decreasing gating-spring tensions, opening and closing the ion channels, resulting in an oscillating receptor current. We hypothesize that decreasing extracellular calcium, decreases the heights of the shorter stereocilia, increasing resting gating-spring tensions, which increases the resting receptor current and decreases the bundle’s resting deflection. To determine the plausibility of this mechanism, we build a mathematical model of an outer-hair-cell bundle and calibrate the model using seven independent experimental observations. The calibrated model shows that the mechanism is quantitatively plausible and predicts that a decrease of only 10 nm in the heights of the shorter stereocilia when extracellular calcium is lowered is sufficient to explain the observed increase in the resting receptor current. The model predicts the values of nine parameters and makes several additional predictions.

## Introduction

In our ears, outer-hair-cell bundles (OHBs) convert sound-induced forces into receptor currents [1, 2]. These receptor currents drive a process known as the cochlear amplifier, which is responsible of our hearing’s high sensitivity, sharp frequency selectivity, and wide dynamic range. Although OHBs are required for normal hearing, we do not fully understand how they work. Experiments show that the OHB’s resting receptor current (no stimulus forces) increases when the extracellular calcium concentration is lowered to physiological levels [3–6]. We propose a mechanism that can account for changes in an OHB’s resting state owing to changes in extracellular calcium.

An OHB comprises filaments, known as stereocilia, emanating from the outer hair cell’s apical surface (Fig 1) [1, 2]. Within an OHB, stereocilia of similar height form rows and stereocilia of differing height form columns. In a column, stereocilia from different rows are linked by gating springs, made of proteinaceous tip links and other elements in series with the tip links. At the lower ends of the gating springs sit ion channels, embedded in the shorter stereocilia of rows 2 and 3. Stimulus forces toward the tallest row (row 1) deflect the stereocilia, which pivot at their insertion points, increase gating-spring tensions, and open the ion channels, through which the receptor current flows. The receptor-current response to stimulation depends on the resting current, which in turn depends on the extracellular calcium concentration [3–6].

**Fig 1.**
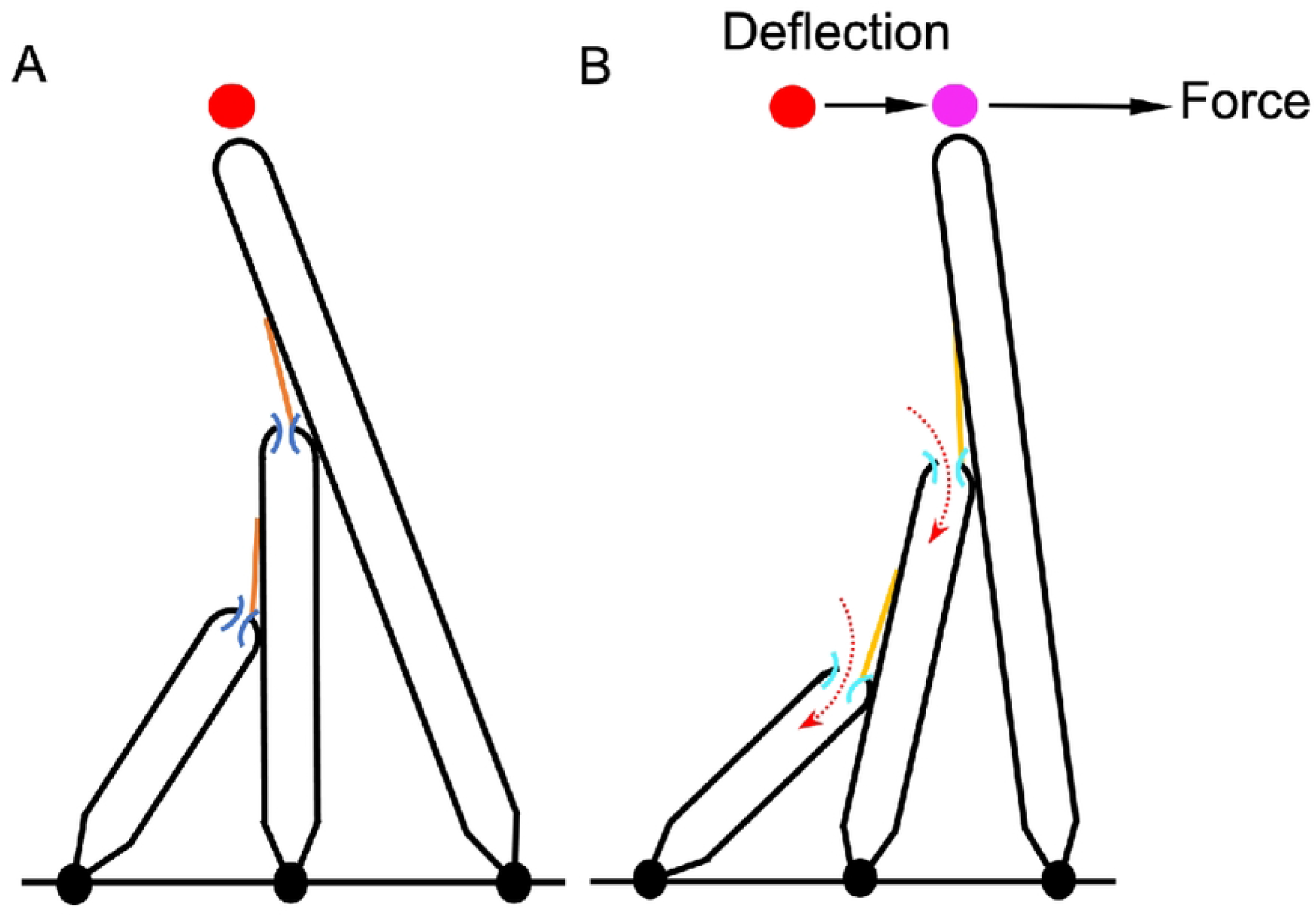
OHB morphology and response to stimulation. (A) A schematic of the OHB is shown. An OHB consists of three rows of stereocilia decreasing in height from row 1 (tallest) to row 3 (shortest). Gating springs (orange) link each stereocilium with its shorter neighbor. The stereocilia in row 2 and row 3 have ion channels (blue) at their tips. (B) A schematic of the OHB responding to a stimulus force is shown. When an OHB is deflected towards row 1 (magenta colored dot) in response to a stimulus, the gating springs extend (light orange) and the channels open (light blue), increasing the ionic currents into the OHB (dashed red arrows). The sum of the currents is called the receptor current.

The extracellular calcium concentration is low (*<* 50 *μ*M) under normal physiological conditions [7]. In low extracellular calcium, the resting current is about half the maximum receptor current and is a major component of inner ear’s silent current [7, 8]. However, in higher calcium concentration environments (*≥* 0.5 mM), the resting current is smaller (*<* 0.2 of the maximum receptor current) [7]. In other words, decreasing extracellular calcium opens the ion channels. Decreasing the extracellular calcium concentration also decreases the heights of the shorter stereocilia, owing to the change in calcium entering through the open ion channels at rest [9]. We propose that lowering calcium increases the resting current by decreasing the heights of the shorter stereocilia.

Here, we quantitatively evaluate the plausibility of our proposal. We build a mathematical model of the OHB, calibrated using many experimental observations. The calibration procedure predicts the values of several OHB parameters. Using the model, we show that decreasing stereocilium heights in low calcium can explain the increase in the resting current. The mathematical model also predicts that the resting OHB displacement and resting gating-spring tensions change when the calcium concentration is changed.

## Methods

The OHB mathematical model comprises 28 independent and identical columns of stereocilia. Each column contains three stereocilia of differing height linked by gating springs (Fig 1). The stereocilia remain in sliding contact with their neighbor in each column when they pivot [10]. Consequently, the displacements of the shorter stereocilia (rows 2 and 3) and the gating lengths (gating length equals the stereocilium radius plus the tip-link length) are dictated by the displacement of the tallest row of stereocilia (row 1) and the morphology of the OHB (Fig S1 and Table S1 in S1 Text). The stereocilia pivot in 2D and have the same pivot stiffnesses and unloaded states. Likewise, the gating springs have same stiffnesses and unloaded lengths. The resting gating length equals the stereocilium radius plus the measured resting tip-link length.

Each gating spring is attached to an MET channel with two states, open and closed [11]. The current through the channel normalized by the maximum current equals the open probability of the channel, which depends on the gating length. Opening a channel decreases the gating-spring length by an amount known as the gating swing [12]. The gating-spring length equals the gating length minus the product of the gating swing and the channel open probability. The receptor current equals the sum of the currents through rows 2 and 3. We normalize the receptor current using the maximum receptor current.

The number of stereocilia, the spacing between the stereocilium pivot points, the widths of the stereocilia, and the heights of the stereocilia are based on published experimental observations on OHBs from the 4-kHz characteristic-frequency region in rats (Table S1 in S1 Text) [13]. Using published experimental observations, we create constraint equations that enable us to derive nine additional parameter values describing the OHB model. This OHB model and the fitting procedure are described in detail in S1 Text.

Mathematica 13 was used to solve the mathematical model. Data from Johnson et. al (2011) was extracted using WebPlotDigitizer (automeris.io) and fitted using Mathematica 13 [4].

## Results

To determine how the resting state of the OHB depends on extracellular calcium, we build a model of the OHB constrained by published experimental observations. The morphology of the OHB model (stereocilium number, heights, widths, and pivot separations) in high calcium is based on published experimental observations (Table S1 in S1 Text). The OHB is described by identical columns of three stereocilia and these stereocilia remain in sliding contact. These choices greatly reduce the number of parameters and variables needed to describe the OHB, which enables us to calculate the values of all the remaining OHB parameters (Methods and S1 Text).

### The OHB model quantitatively accounts for seven independent experimental observations

In response to a stimulus toward row 1, the stereocilia are displaced in the positive direction and the MET channels open. The model OHB reproduces the measured receptor-current dependence on the OHB’s displacement (the tip displacement of the row-1 stereocilium), including the resting receptor current in high calcium (Fig 2A and Table 1). When the gating springs are cut in experiment, the OHB’s stiffness (slope of the force versus displacement curve) decreases (Fig 2B,C and Table 1). The model OHB captures the measured stiffnesses of the OHB with (in high calcium) and without gating-springs. For low-calcium, the resting length of the tip link and the resting receptor current in the OHB model equal the values measured experimentally (Table 1). When the gating springs are cut in low calcium, the resting displacement of the OHB increases in experiment, a change which is also matched by the OHB model (Fig 2B–D and Table 1). Overall, the model OHB quantitatively fits seven independent experimental observations.

**Fig 2.**
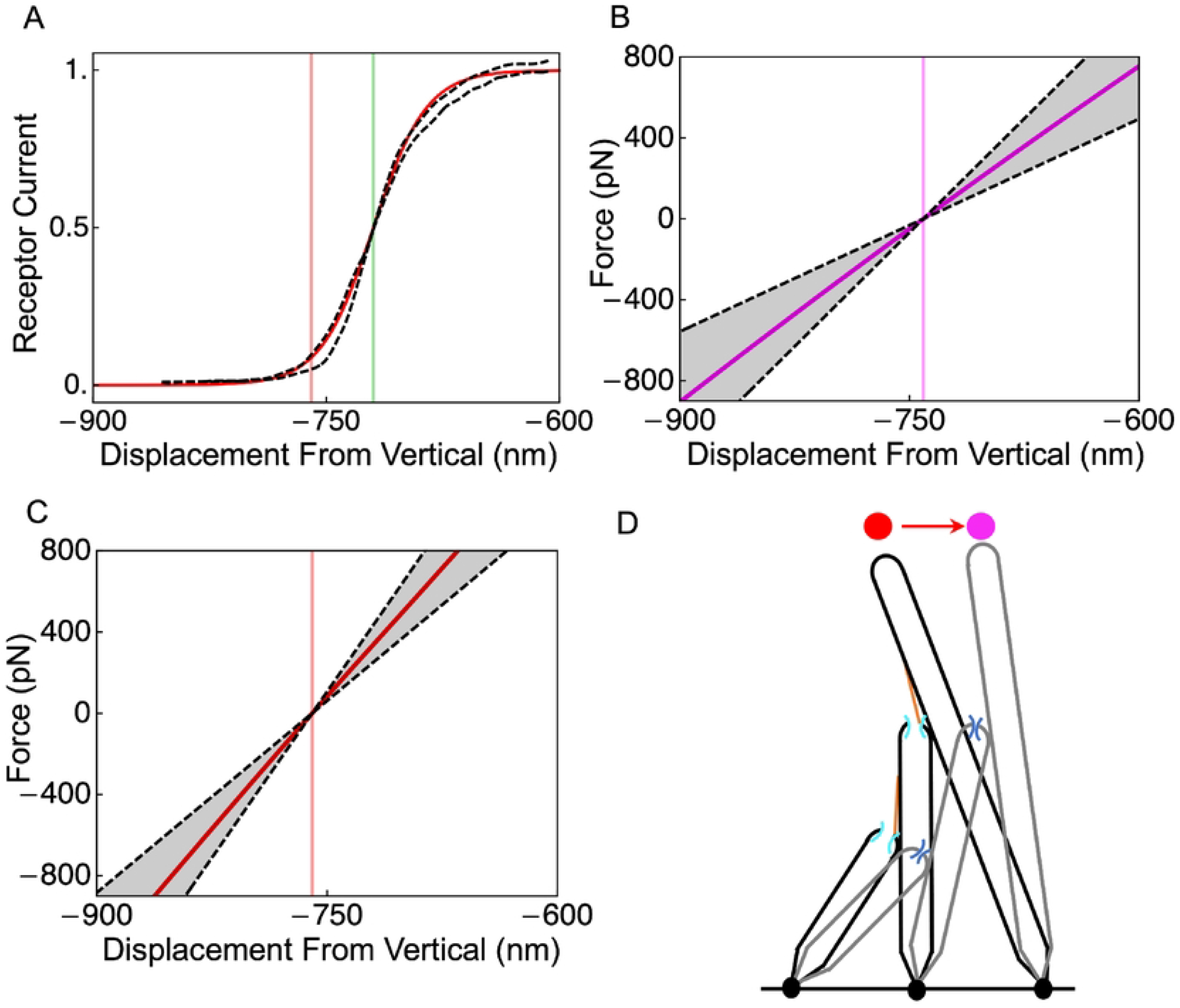
The OHB model fits published experimental observations. (A) The receptor current is shown versus the OHB displacement (known as the activation curve) for the model (red line) and for experiment (black dashed lines). The model agrees with the average of the forward and backward cycles of stimulation in the experiment. The extracellular calcium concentration is high. Vertical lines indicate OHB displacement at rest (red) and at half-activation (green) for the model. (B) The force applied to the OHB versus its displacement is shown for the model (pink line) and for experiment (black dashed lines indicate the mean +/-the standard deviation) when the OHB has no gating springs. The vertical pink line indicates the resting OHB displacement for the model. (C) The force applied to the OHB versus its displacement is shown for the model (red line) and for experiment (black dashed lines indicate the mean +/-the standard deviation) when the OHB has gating springs. The vertical red line indicates the resting OHB displacement for the model. (D) Schematic representations of the OHB are shown with gating springs (black) and without gating springs (grey), illustrating that the resting OHB displacement increases (red arrow) when the gating springs are broken. The dot indicate the resting displacement with (red) and without (pink) gating springs.

**Table 1.** Emergent property values (mean *±* standard deviation). Seven independent experimental observations are listed. The resting current in high calcium 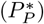 is determined by the measured activation curve width (2Δ_*J*_) and center (*X*_*J*_).

| Emergent property | Experiment | Model |
| --- | --- | --- |
| **Low calcium** |  |  |
| Resting current ($P_E^*$) | 0.5 [4, 14] | 0.5 |
| Resting tip-link length ($x_{TL}^*$) | $186 \pm 38$ nm [15] | 186 nm |
| OHB stiffness without gating springs ($K_{HB}^{noGS}$) | $5.5 \pm 2$ mN/m [14] | 5.5 mN/m |
| Deflection from breaking gating springs ($\Delta_E$) | $45 \pm 10$ nm [14] | 45 nm |
| **High calcium** |  |  |
| Activation-curve width (0.27–0.73) ($2\Delta_J$) | 34 nm [4] | 34.1 nm |
| Activation-curve center relative to rest ($X_J$) | 40 nm [4] | 40 nm |
| Resting current ($P_P^*$) | $0.087 = (1 + \exp(X_J/\Delta_J))^{-1}$ [4] | 0.087 |
| OHB stiffness with gating springs ($K_{HB}$) | $8.6 \pm 2.3$ mN/m [14] | 8.6 mN/m |

### The OHB model predicts the values of nine parameters

The model OHB predicts the values of 17 unknowns (9 parameter values and 8 state values) based on 17 constraints and prior experimental observations (Fig 3, Tables 1 and 2, S1 Text, Table S2 in S1 Text). Following prior work, we assume that myosin motors set the resting gating-spring lengths to be the same [13]. Then, based on the measured resting receptor current in low calcium of 0.5, the resting currents in low calcium are 0.5 for all the MET channels. The sliding contact assumption enables us to find the resting displacement of the OHB when its gating springs are cut and then to find the resting displacement of the OHB with intact gating springs (−786 nm) from the measured displacement of the OHB caused by cutting the gating springs. The resting displacements are negative relative to the vertical owing to the balance of forces and torques at rest.

Combining the resting displacement of the intact OHB with its geometry yields the resting sliding contact points. In the model OHB, the resting gating lengths equal the stereocilium radius plus the measured resting length of the tip links. We calculate the upper tip-link attachment locations using the resting gating lengths and the resting sliding contact points. The gating length required for the channel current to be 0.5 (the half-activation length) equals the resting gating lengths in low calcium. Using the measured stiffness of the OHB after its gating spring have been cut, we find a pivot stiffness value of 1.0 fN.m/rad, which is smaller than prior work using an identical-gating model (1.3 fN.m/rad), because the prior model is simpler and we use more stereocilia (84 here versus 70.6 in prior work) [14].

**Fig 3.**
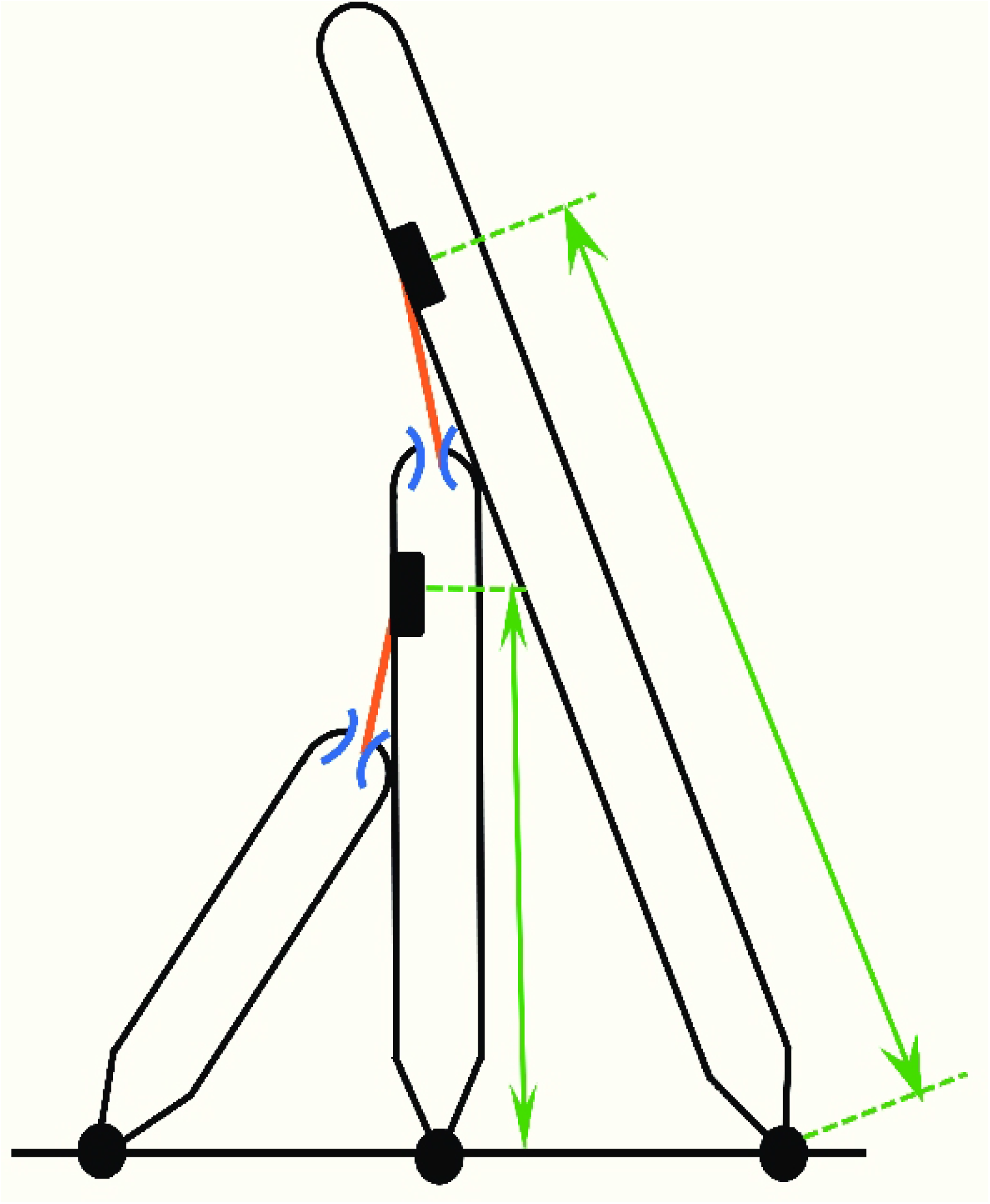
The OHB model predicts the values of several OHB properties. A schematic representing the OHB is shown highlighting the parameters deduced using the OHB model (values are summarized in Table 3). These parameters are the upper tip-link insertion distances for row 1 and row 2 (green arrows; *b*_1_ and *b*_2_), the stiffness of the stereocilium pivots (black dots; *κ*), the stiffness (*k*_*gs*_) and the unloaded length (*x*_*u*_) of the gating springs (orange), the gating swing of the channels (blue; *d*), and the half-activation length of the gating springs (orange/blue; the half-activation length depends on the gating springs and the channels; *x*_*h*_).

Given the calculated values of the upper tip-link attachment points, the half-activation length, and the pivot stiffness we find the gating-spring stiffness (2.9 mN/m), the unloaded length of the gating spring (319 nm), the gating swing (0.6 nm), and the heights of the row 2 (2110 nm) and 3 (1310 nm) stereocilia in high calcium using the measured stiffness of the OHB in high calcium, the measured OHB activation curve in high calcium, and the measured displacement of the OHB caused by cutting the gating springs in low calcium. The predicted value of the gating-spring stiffness (2.9 mN/m) is smaller than the value found using an identical-gating model (3.7 mN/m) in prior work, because the identical-columns model is simpler, we use fewer gating springs (56 here versus 65 in prior work), and we use larger pivot spacings (603 nm on average here versus 462 nm in prior work) [14]. In contrast, the predicted value of the gating swing equals the value assumed in a prior OHB model [16].

**Table 2.** Predicted parameter values and resting-state variable values. The parameters are the upper tip-link insertion distances for row 1 and row 2 (*b*_1_ and *b*_2_), the stiffness of the stereocilium pivots (*κ*), the stiffness (*k*_*gs*_) and the unloaded length (*x*_*u*_) of the gating springs, the gating swing of the channels (*d*), the half-activation length of the gating springs (*x*_*h*_), and the heights of row 2 and row 3 in high calcium (*l*_2,*P*_ and *l*_3,*P*_). The resting-state variables are the resting OHB displacement without gating springs 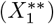, the resting OHB displacement in low calcium 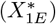, the resting gating lengths in low calcium 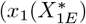 and 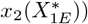, the resting tip-link length plus the stereocilium radius in low calcium 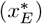, the resting OHB displacement in high calcium 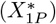, and the resting gating lengths in high calcium 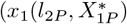 and 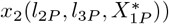.

| Unknown | Prediction | Unknown | Prediction |
| --- | --- | --- | --- |
| Low calcium |  | High calcium |  |
| Parameter | Value | Parameter | Value |
| $\kappa$ | 1.0 fN.m/rad | $l_{2,P}$ | 2110.0 nm |
| $b_1$ | 2490.1 nm | $l_{3,P}$ | 1309.7 nm |
| $b_2$ | 1589.3 nm | $k_{gs}$ | 2.9 mN/m |
| $x_h$ | 330.5 nm | $d$ | 0.6 nm |
| $x_u$ | 319.1 nm | | |
| Variable | Value | Variable | Value |
| $X_1^{**}$ | -741.0 nm | $X_{1P}^*$ | -759.8 nm |
| $X_{1E}^*$ | -786.0 nm | $x_1(l_{2P}, X_{1P}^*)$ | 324.9 nm |
| $x_1(X_{1E}^*)$ | 330.5 nm | $x_2(l_{2P}, l_{3P}, X_{1P}^*)$ | 324.9 nm |
| $x_2(X_{1E}^*)$ | 330.5 nm | | |
| $x_E^*$ | 330.5 nm | | |

### Increasing the heights of rows 2 and 3 explains the change in resting state from low to high calcium

To account for the measured differences in resting receptor current (0.087 in high calcium versus 0.5 in low calcium), we allow the heights of row 2 and 3 to differ in high and low calcium in the model OHB (Fig 4). The heights in low calcium are based on optical and electron microscopy as described previously [13]. The OHB model predicts that an increase in height of about 10 nm in high calcium decreases the resting tensions (31.9 pN in low calcium to 16.6 pN in high calcium) and increases the resting displacement of the OHB (−786 nm in low calcium to -760 nm in high calcium). Similar height increases are predicted for row 2 and row 3.

**Fig 4.**
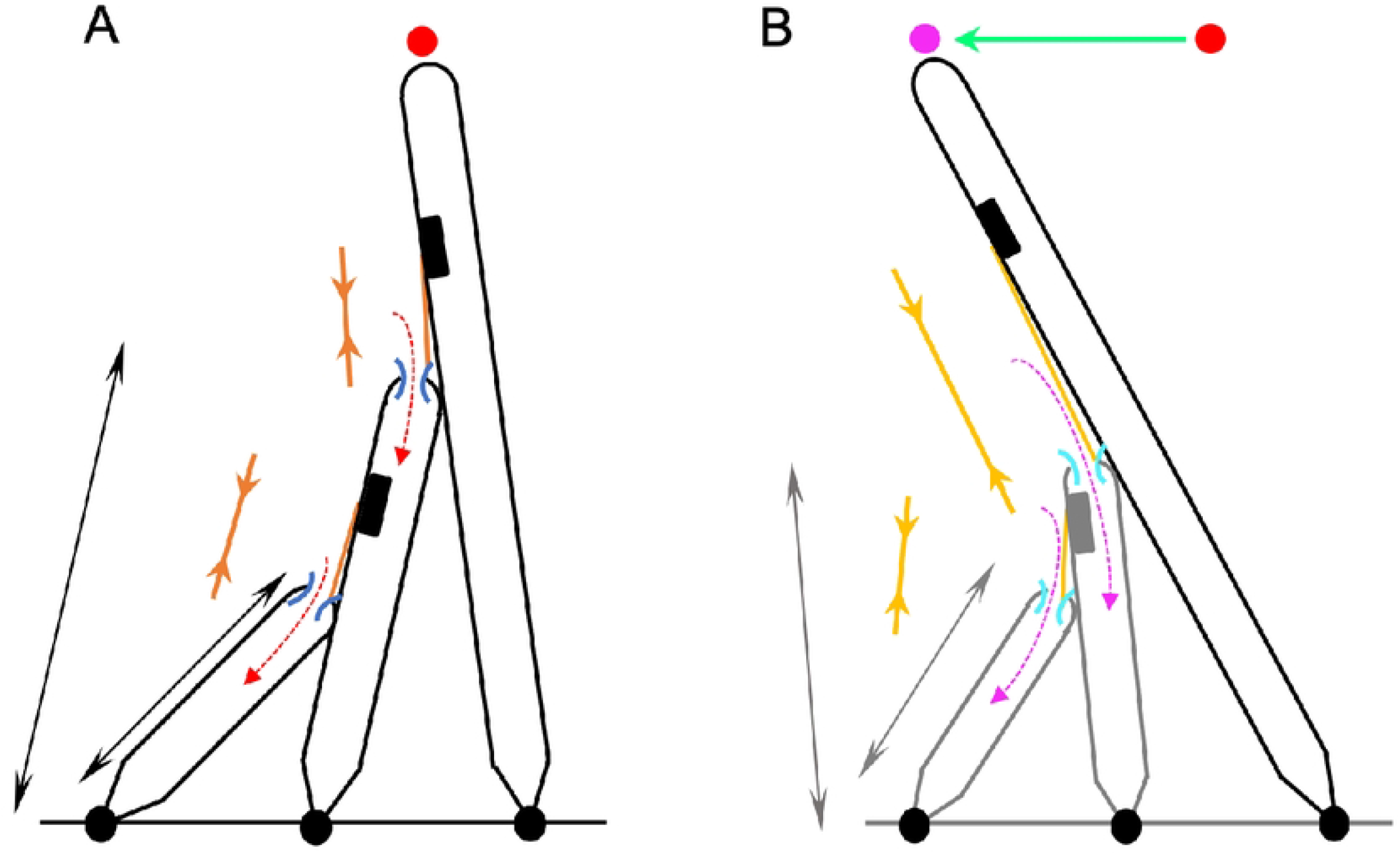
A decrease in the row-2 and row-3 heights in low extracellular calcium increases the resting receptor current and decreases the resting deflection of the OHB. (A) A schematic representing the OHB in high extracellular calcium is shown. The resting current (red-dashed arrows), resting channels (dark-blue arcs), resting deflection from the vertical (red dot), resting tensions (dark-orange arrows), and the heights of rows 2 and 3 (black arrows) are indicated. (B) A schematic representing the OHB in low extracellular calcium is shown. The resting current (pink-dashed arrows, increased relative to high extracellular calcium), resting channels (light-blue arcs, more open relative to high extracellular calcium), resting deflection from vertical (pink dot, the green arrow indicates the decrease relative to high extracellular calcium), resting tensions (light-yellow arrows, increased relative to high extracellular calcium), and the row-2 and row-3 heights (grey, decreased relative to high extracellular calcium) are indicated.

We fit the model OHB to the receptor-current activation curve and OHB stiffness near the resting state in high calcium (Table 1, Fig 2). The model OHB predicts that the activation curve center decreases in low calcium by 66 nm (Fig 5A). Owing to the geometry of an OHB column, the model OHB also predicts that the OHB’s stiffness decreases with OHB displacement in high and low calcium (Fig 5B). However, the angular displacements of the stereocilia increase almost linearly and their sliding contact positions decrease almost linearly with the OHB’s displacement (Figs S2 and S3 in S1 Text). In high and low calcium, MET channel gating decreases the OHB’s stiffness when the receptor current is near 0.5, a phenomenon known as gating compliance [12]. Within the physiological range of OHB displacements, the row-2 and row-3 gating lengths are very similar, which make the channel currents versus the OHB’s displacement almost identical (Figs. S4 and S5 in S1 Text).

**Fig 5.**
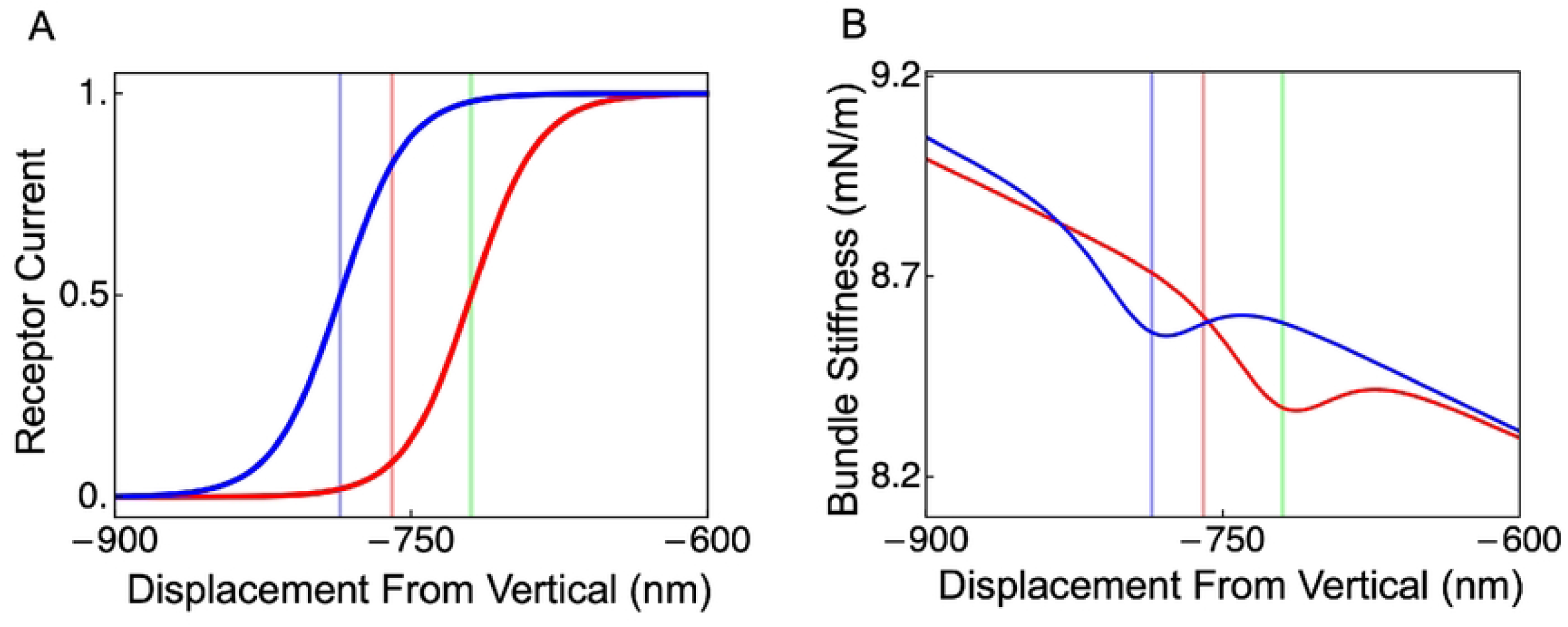
The OHB model predicts the activation curve in low extracellular calcium and the OHB stiffnesses in high and low extracellular calcium. The receptor currents (A) and the OHB stiffnesses (B) are shown versus the OHB displacement in high (red line) and low (blue line) extracellular calcium. Vertical lines indicate OHB displacements in high extracellular calcium at rest (red), in high extracellular calcium at half activation (green), and in low extracellular calcium at rest (blue).

## Discussion

Our results show that shorter-row height changes owing to changes in calcium concentration is a plausible mechanism for explaining resting-state changes in OHBs. Accounting for seven different experimental observations from three different papers using a single mathematical model, suggests that the experimental observations are quantitatively consistent across these publications [4, 14, 15]. In addition to predicting the values of nine parameters, the mathematical model makes several other predictions.

The predicted increase of 5.6 nm in the resting gating length (equaling the resting tip-link length plus the stereocilium radius) caused by lowering the calcium concentration is consistent with the measured increase of 21 *±* 69 nm in the resting tip-link length (164.4 *±* 57.3 nm in 1 mM calcium and 185.8 *±* 37.6 nm in 50 *μ*M calcium; mean *±* standard error) [15]. This agreement implies that the stereocilium radius need not change to account for the predicted gating-length change.

When extracellular calcium is decreased, the mathematical model predicts that the resting OHB displacement decreases by 26 nm, the activation curve shifts in the negative direction, and the center of the nonlinearity associated with channel gating shifts in the negative direction (Table 2; Fig 5). The predicted 26 nm increase in the resting OHB displacement when the calcium concentration is increased is similar to the measured increase in displacement of about 20 nm when the calcium concentration is increased from about 0 mM to 0.5 mM [14]. Negative shifts in the activation curve and center of the nonlinearity relative to the resting state have been observed [3–5, 17]. A prior mathematical model of receptor-current adaptation (processes that maintain the sensitivity of a bundle subjected to a static displacement stimulus) also predicts that activation curves shift in the negative direction when extracellular calcium is lowered, suggesting that there may be several contributions to the activation curve shift [17].

The negative resting displacement of the OHB predicted by the model in high calcium (−760 nm; arcsin(*−*760*/*4100) = *−*11^*°*^ from the vertical, in which the OHB height is 4100 nm; Table 2 and Table S1 in S1 Text) is quantitatively consistent with prior observations of OHBs (*−*8^*°*^ to *−*12^*°*^ on average) [18]. We find a similar negative resting displacement in low calcium *−*786 nm (Table 2). However, the OHB is embedded in an overlying membrane, called the tectorial membrane *in vivo* [1, 2]. *In situ* measurements in high calcium show positive resting displacements, implying that the tectorial membrane biases the OHB in the positive direction (15^*°*^ on average at the cochlear apex) [19]. Importantly, recent *in vivo* measurements suggest that the tectorial membrane does not bias the resting OHB displacement [7]. The resting OHB receptor current in low calcium is about 0.5 with and without the tectorial membrane and depends on the resting displacement. It is unlikely that receptor-current adaptation can maintain a receptor current close to 0.5 if the resting displacement were shifted from our calculated value of -786 nm to 1161 nm (4100 *×* sin(15^*°*^) nm; Table 2 and Table S1 in S1 Text) by the tectorial membrane, because there is little adaptation for OHB displacements larger than 500 nm [20–22]. Determining whether the resting displacement is positive or negative *in vivo* is important for our understanding of the cochlear amplifier [23, 24].

Unlike a prior 3D mathematical model of the OHB, the essentially 2D model (all columns are identical and independent and are described by the 2D motions of a single column) presented here produces similar channel currents in rows 2 and 3 (Fig S5) [13]. The current similarity is a consequence of the 2D and sliding-contact approximations, because a 3D OHB model lacking the sliding-contact assumption shows that row-2 and row-3 currents differ substantially [13]. However, these approximations enable us to better understand the OHB and to fit the model exactly to the experimental data (Table 1 and Table S2 in S1 Text). We expect the 2D model predictions to hold qualitatively in 3D, because 3D models are extensions of 2D models.

OHB stiffness has been observed to increase when the calcium concentration is lowered from 1.5 mM to 0.02 mM [17]. We find that this change in OHB stiffness is not explained by changing the row-2 and row-3 stereocilium heights, implying that additional calcium-dependent mechanisms are needed to explain the observation (Fig 5). Possible mechanisms to explain the OHB stiffness increase when calcium decreases include, decreasing the unloaded length of the gating spring, increasing gating-spring stiffness, and increasing the stiffness of an element in series with the gating spring [17, 25].

Why might extracellular calcium affect stereocilium heights? One possibility is that a decrease in extracellular calcium decreases calcium influx through the channels, decreasing the rate of actin polymerization in row-2 and row-3 stereocilia, decreasing row-2 and row-3 heights [9]. However, the rate of height changes owing to actin polymerization is limited. The polymerization rate is 1-10 monomers per second and each actin monomer is about 6 nm long [26]. To increase stereocilium heights by 10 nm would require 1.2 monomers on average, which would take 0.12-1.2 seconds. Another possibility, is that there are calcium-dependent elastic elements within stereocilia, whose resting lengths regulates the resting stereocilium heights. Decreasing extracellular calcium decreases calcium influx, which might increase the stiffness of the elements, decreasing stereocilium heights. An element within OHB stereocilia with a calcium-dependent stiffness has been proposed previously to affect the resting current, but the potential effects on stereocilium heights was not described [25]. The rate of height changes owing to this mechanism is unknown.

Several calcium-dependent process might change the resting state of the OHB [27]. Our proposal is independent from some of these processes (e.g., lipid bilayer modulation by calcium), but might contribute to other processes. Like decreasing extracellular calcium, depolarization to a positive membrane potential decreases intracellular calcium and increases the resting current magnitude, suggesting that depolarization might decrease row-2 and row-3 heights [5, 6, 9]. Depolarization affects the receptor current on timescales of 100 ms to several seconds, so an actin polymerization mechanism would be sufficiently fast to contribute to depolarization regulation of the resting current [5, 6]. In contrast, slow adaptation is a calcium-dependent process that decreases the receptor current with a timescale of about 20 ms in OHBs, implying that height changes based on actin polymerization alone are not sufficiently rapid to account for slow adaptation [5, 27]. However, a calcium-dependent elastic element mechanism that changes stereocilium heights might be sufficiently fast to contribute to slow adaptation [25]. Finally, it is possible that calcium-dependent height changes are smaller than we predict, because other calcium-dependent processes might account for some of the resting-current changes [27].

## Conclusions

We propose a new mechanism for regulating the resting state of the OHB that brings together several different experimental observations. Decreasing extracellular calcium decreases the heights of rows 2 and 3, increasing resting gating-spring tensions, deflecting the OHB in the negative direction and increasing the resting receptor current. The mathematical model predicts the values of nine parameters and the resting values of eight variables. Some predictions agree with prior experimental data, some predictions imply that additional mechanisms are needed to explain experimental data, and all predictions can in principle be tested experimentally.

## Acknowledgments

This work was funded by Stanford Otolaryngology — Head & Neck Surgery, the Stanford Initiative to Cure Hearing Loss, and a Postdoctoral Support Award for R. C. from the Maternal and Child Health Research Institute at Stanford.

## Supporting information

**S1 Text. chatterjeeSI.pdf** Mathematical model description and supporting figures.

**S2 Code. resting_ca_dependence.nb** Mathematical model code (Mathematica 13.3.1).

**S3 Data. Jd1e1b.csv** Experimental data used for fitting mathematical model.

**S4 Data. Jd1e2b.csv** Experimental data used for fitting mathematical model.

**S5 Data. Johnsonsine.csv** Experimental data used for fitting mathematical model.

**S6 Data. Johnson20111E.csv** Experimental data used for fitting mathematical model.

## Notes

### Competing Interest Statement

The authors have declared no competing interest.

